# Comprehensive analysis of pathways in Coronavirus 2019 (COVID-19) using an unsupervised machine learning method

**DOI:** 10.1101/2022.05.18.492441

**Authors:** Golnaz Taheri, Mahnaz Habibi

**Affiliations:** Department of Electrical Engineering and Computer Science, KTH Royal Institute of Technology, Stockholm, Sweden; Science for Life Laboratory, Stockholm, Sweden; Department of Mathematics, Qazvin Branch, Islamic Azad University, Qazvin, Iran

**Keywords:** Coronavirus disease 2019, SARS-CoV-2, Machine learning, Unsupervised learning

## Abstract

The World Health Organization (WHO) introduced “Coronavirus disease 19” or “COVID-19” as a novel coronavirus in March 2020. Severe acute respiratory syndrome coronavirus 2 (SARS-CoV-2) requires the fast discovery of effective treatments to fight this worldwide crisis. Artificial intelligence and bioinformatics analysis pipelines can assist with finding biomarkers, explanations, and cures. Artificial intelligence and machine learning methods provide powerful infrastructures for interpreting and understanding the available data. On the other hand, pathway enrichment analysis, as a dominant tool, could help researchers discover potential key targets present in biological pathways of host cells that are targeted by SARS-CoV-2. In this work, we propose a two-stage machine learning approach for pathway analysis. During the first stage, four informative gene sets that can represent important COVID-19 related pathways are selected. These “representative genes” are associated with the COVID-19 pathology. Then, two distinctive networks were constructed for COVID-19 related signaling and disease pathways. In the second stage, the pathways of each network are ranked with respect to some unsupervised scorning method based on our defined informative features. Finally, we present a comprehensive analysis of the top important pathways in both networks. Materials and implementations are available at: https://github.com/MahnazHabibi/Pathway.

## 1. Introduction

Coronavirus (CoV) is a large family of respiratory viruses that can cause a variety of illnesses ranging from mild to severe diseases including the common cold, to severe acute respiratory syndrome (SARS) [1]. CoV is found in different kinds of animals, including camels, pangolins, and bats [2]. Some of these viruses can evolve and infect humans on rare occasions. The severe acute respiratory syndrome coronavirus 2 (SARS-CoV-2) is a newly evolved virus from this family. Pneumonia caused by the SARS-CoV-2 was introduced by the World Health Organization (WHO) as coronavirus disease (COVID-19).

Even though there are over thousands of clinical trials, there are no approved medications for the COVID-19 [3]. Although SARS-CoV-2 has a lower mutation rate than other coronaviruses, at the same time, genomic diversity is seen both among individual patients and within the same virus class [4, 5]. Genetic diversity allows viruses to adapt to a variety of hosts and circumstances within those hosts. This diversity is mostly associated with disease development, drug resistance, and treatment outcomes. As a result, even minor but continual virus alterations and mutations would reduce the efficiency of vaccines or commonly used drugs for COVID-19 treatments. Therefore, gathering information about the virus’s development and pathology will be critical to controlling the pandemic.

Many researchers are sharing their findings to learn more about SARS-CoV-2’s genome and evolution in the world. Several researchers are focusing on finding a treatment with the help of existing drugs with the drug repurposing method as a faster and less expensive technique [6]. Pathway analysis is a useful method for drug repurposing and understanding different patients’ responses to the virus. Pathway analysis can improve the interpretation of SARS-CoV-2 data by identifying biological pathways of host cells affected by the virus. From a large amount of SARS-CoV-2 related data released, this kind of analysis can help to characterize possible drug targets and drug mechanisms of action. As a result, to find an effective treatment, obtaining knowledge from data that characterizes SARS-CoV-2 host infection is a beneficial approach.

Biological pathways are human representations of a cell’s coordinated molecular operations, which might involve genes, proteins, molecules, tissues, and organs. Signaling pathways, disease pathways, metabolic pathways, and regulatory pathways are the four main groups of these pathways. Generic pathway databases, which include pathways from all groups, are more extensive and widely utilized. From all of the available pathway databases, each database has a distinct number of pathways. However, the pathways of each database only annotate a portion of the genome. When all major pathway databases are combined, most of the pathway databases focus on Homo sapiens-related pathways [7]. Pathway analysis methods can help researchers to identify the role of host genes in altering the biological response to SARS-CoV-2 and different outcomes of COVID-19.

Ranking important and more relevant pathways from all of the COVID-19 associated pathways proposed by recent studies will assist researchers to focus on select sets of genes for further inquiry. Furthermore, observing pathways as functional units of cellular systems may help researchers extract biologically relevant information from microarray data. Therefore, various research groups have been inspired to look at gene sets rather than single genes as a result of these considerations.

In this work, we developed an unsupervised machine learning approach to identify important disease pathways crosstalk between underlying diseases and COVID-19. We also tried to recognize important signaling pathways of the host protein which could help to identify effective COVID-19 treatment. For this purpose, we constructed two biological networks corresponding to the COVID-19 related signaling and disease pathways. Then, we defined six informative structural features for each pathway as a vertex in the network. Finally, we calculated the Laplacian Score with respect to our predefined features for each pathway and introduced the high score pathways with meaningful relationships to COVID-19 as candidate pathways for more investigation.

## 2. Preliminaries

Groups of proteins that interact in the body are referred to as ‘‘cellular processes”. To understand the cell, we must understand how these spatially and temporally organized interactions lead to biological processes [8]. The conventional approach to studying cellular function has improved our understanding of disease, infection, drug development, and evolution [9]. In this work, we define data and networks as “molecular” if they are concerned with interactions between particular biological molecules. There are different approaches for analyzing biological processes using molecular interaction networks. Protein-protein interaction (PPI) data is often used to create networks in which proteins are shown interacting with other functionally associated proteins. This results in the formation of “functional modules”, which are subnetworks that share a common function [10]. In PPI networks, the edges connect each protein to all of its other interacting proteins. Proteins may participate in various functions depending on the interactions they create in different cellular contexts and subcellular components [11]. In this work, we introduce a representation of cellular functions that uses pathways rather than genes. The pathway includes a set of proteins (and complexes) that interact with each other serially and create signaling, metabolic, and disease pathways [7]. This concept allows us to group sets of proteins known to interact under certain conditions. In this study, we did not use gene expression data and assumed that gene expression represents protein levels. The pathway model allows proteins to be characterized independently in multiple pathways.

Among many types of biological pathways, those that are involved in metabolism, regulation of genes, and the transmission of signals are well-known [12]. The chemical reactions that take place in our bodies are mediated by metabolic pathways. Some other metabolic pathways mostly help to produce molecules. Regulatory pathways turn genes on and off. This is such an important task because genes supply the formula by which cells produce proteins, and proteins are the key elements needed to carry out almost every task in the human body. Signaling pathways move a signal from the outside of the cells into the inside of them. Different cells are capable of receiving specific signals via structures on their surface called receptors. After interacting with these receptors, the signal travels into the cell. The message of a signal is transmitted by some kinds of spatial proteins that start a specific reaction in the cell. For example, a signal from outside the cell could lead the cell to create a particular protein inside the cell. After that, the produced protein may be a signal that causes the cell to move [12]. Figure 1 shows a schematic view of a biological pathway as a series of interactions between different molecules in a cell that leads to a specified product or a change in a cell. As we mentioned earlier, the most common biological pathways are involved in metabolism, the regulation of gene expression, and the transmission of signals.

**Figure 1:**
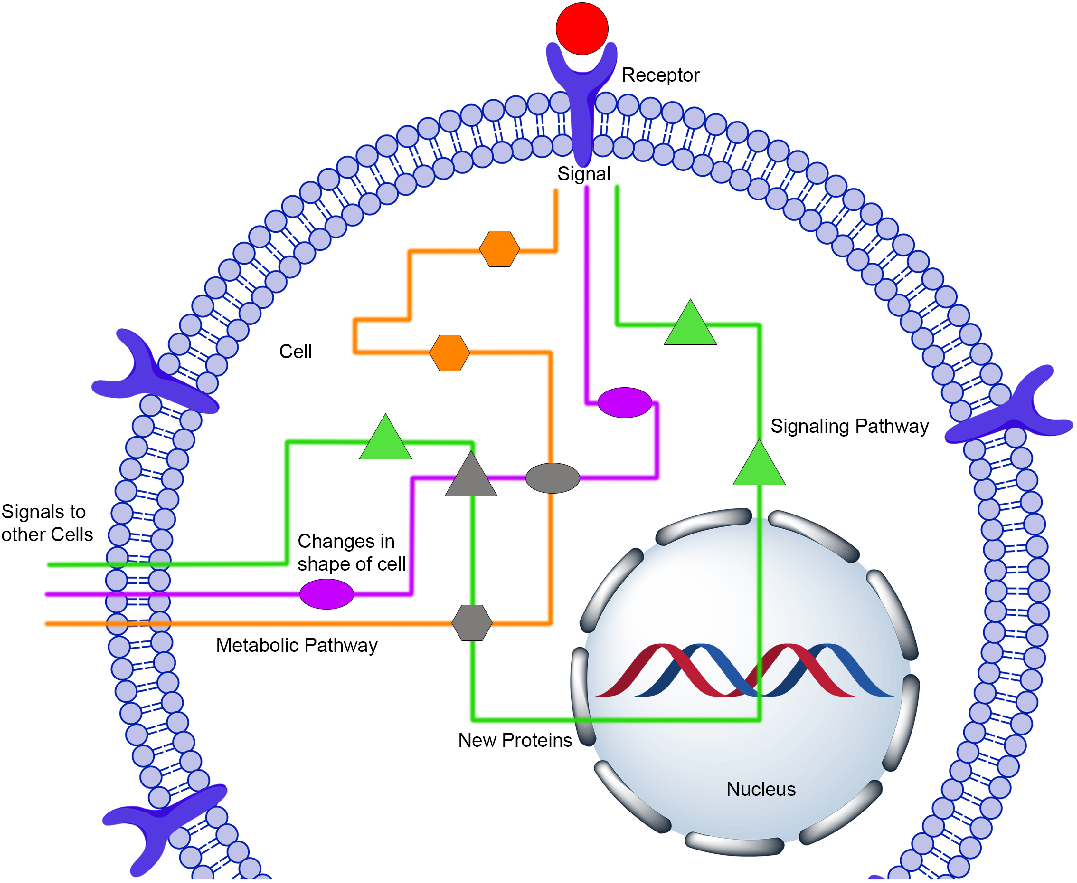
A schematic view of a biological pathway.

In the pathway analysis methods, the position of a given gene in pathways determines its function in some way [13]. Upstream genes can affect the downstream responses of pathways. In addition, genes with a high degree (hubs) in pathways participate in important biological processes that are associated with the diagnosis and therapy of different diseases [14]. For example, p53 is an important upstream gene with a high degree in the P53 signaling pathway, which is the most repeatedly mutated gene in different types of cancer. This gene plays an important role in tumor suppression through transcriptional regulation of its downstream target genes like MYC and EGR1 [15]. Another important upstream gene is RAS. This gene contains several important proteins (KRAS, NRAS, HRAS, RRAS, RRAS2, and MRAS). It has a high degree in the MAPK signaling pathway. The RAS mutation can cause a broad range of genetic disorders [16]. Recently, several methods consider the structure of pathways, such as centrality, betweenness, and weight-based pathway enrichment.

## 3. Related Work

SARS-CoV-2 has become an existing health crisis and threat to the entire world. Thus, the world needs to be fast in recognizing suitable information and medication to restrict the spread of this disease. The large-scale data of COVID-19 patients can be integrated and analyzed by state-of-the-art machine learning algorithms for a better understanding of the pattern of viral spread, and to improve diagnosis speed and accuracy. These algorithms also develop novel, effective therapeutic approaches and could identify the most sensitive people based on personalized genetics. After a short period of time since the COVID-19 outbreak, advanced machine learning techniques have been used in the classification of COVID-19 genomes [17], evaluation of CRISPR-based COVID-19 detection [18], survival prediction of severe COVID-19 patients, and finding potential drug candidates for COVID-19 treatments [19].

Current efforts have been made to develop effective and novel diagnostic approaches with the help of machine learning algorithms. As an example, machine learning-based screening of SARS-CoV-2 evaluation techniques using a CRISPR-based virus detection system was confirmed with high accuracy and speed [18]. Neural network-based classifiers were developed for large-scale screening of COVID-19 patients based on their specific respiratory patterns [20]. Another deep learning-based method for COVID-19 patients’ CT image analysis was constructed for the automated diagnosis and monitoring over time [21]. The fast development of automated diagnostic systems based on artificial intelligence and machine learning methods can increase diagnostic accuracy and speed. These methods also save and protect healthcare workers by decreasing their contact with COVID-19 patients.

In [22] new drug-like compounds for COVID-19 treatment have been developed using a deep learning pipeline. Another deep learning system developed by Google DeepMind, AlphaFold, predicted protein structures associated with COVID-19 [23]. Traditional experimental methods can take months to complete this work. In addition, a new machine learningbased tool proposed COVID-19 vaccine candidates [24]. Data from worldwide COVID-19 treatment centers necessitates advanced machine learning techniques. These methods could be used to evaluate new patients by analyzing their personalized therapeutic effects.

## 4. Material and Methods

As an input, we selected a set of candidate COVID-19 related genes. This candidate set contains most of the genes that have been suggested as drug targets for COVID-19 in the Drug Bank [25]. Then, we extracted the corresponding pathways from Kyoto Encyclopedia of Genes and Genomes dataset (KEGG) for these selected genes [26]. In the first step of this work, we constructed two biological networks corresponding to COVID-19 related signaling and disease pathways. We also defined six informative features for these two networks. In the second step, we used an unsupervised learning method and computed the Laplacian Score, then sorted pathways for each network with respect to the corresponding score. The workflow of the proposed methods to identify important pathways is illustrated in Figure 2.

**Figure 2:**
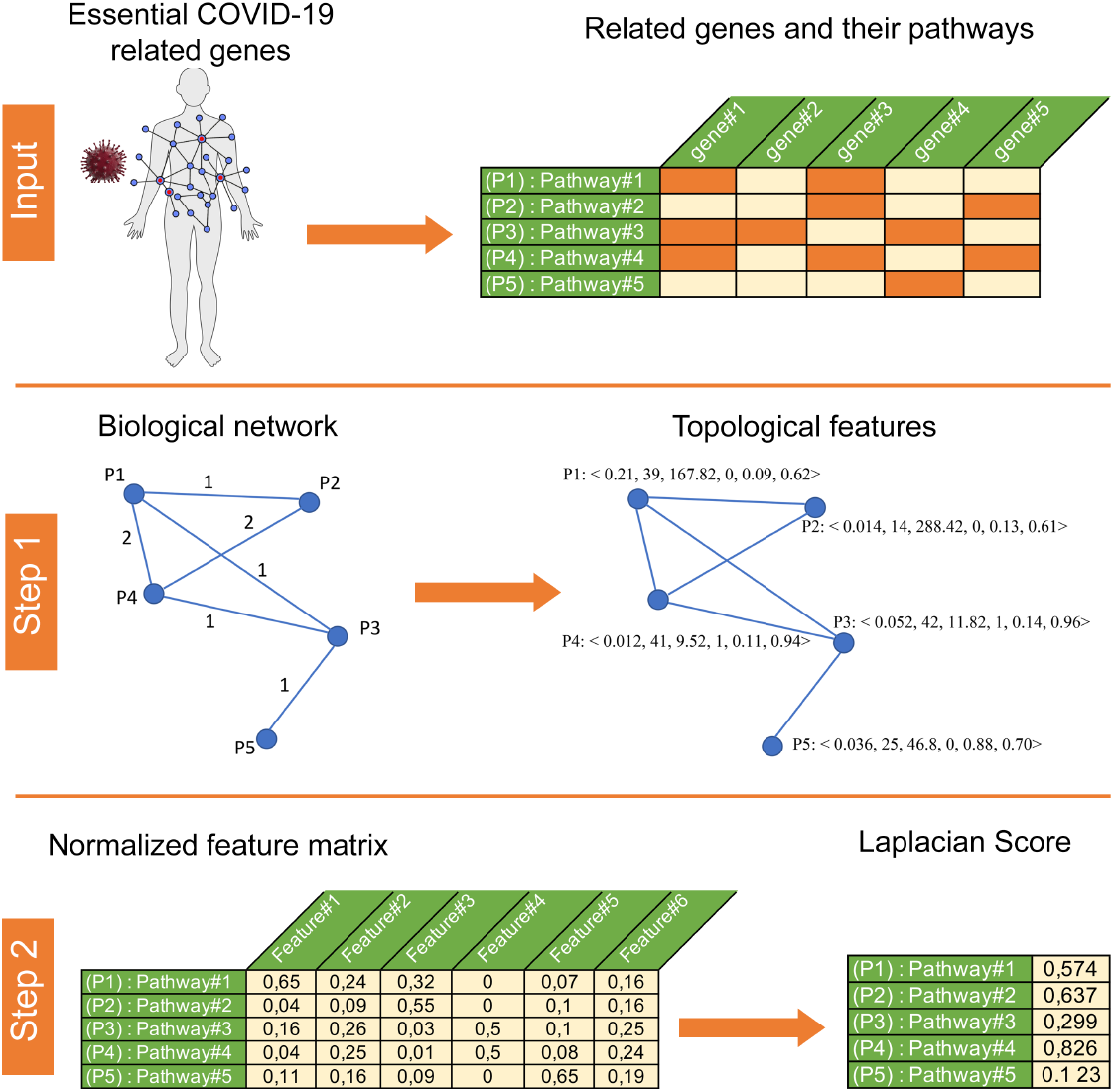
The workflow of the proposed methods.

### A. Protein-protein interaction network and important genes

The physical interaction between two or more protein molecules is called the PPI network. A set of proteins and the physical interactions between them are represented as a proteinprotein interaction network 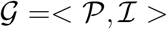. Each vertex in the network represents a protein and each edge, *uv*, in the network shows the interaction between these two proteins, *u* and *v*. When two proteins interact physically, the two vertices, *u* and *v*, are referred to as adjacent. The number of interactions that a vertex, *u*, has with other vertices is called the degree. A set of high-degree vertices is called hub vertices. Hub vertices play an important role and contain important biological information for the cell. Generally, hub vertices and vertices with high centrality measures are selected as important genes in the PPI network. To identify and evaluate the important genes of each COVID-19 related pathway, we will identify the important vertices of the PPI network for all of the genes in that pathway.

### B. Construction of biological network of COVID-19 related pathways

Suppose *S* represents a set of essential genes related to COVID-19, and *P* = {*p*_1_, *p*_2_,…*p_k_*} defines a set of related pathways gathered from the KEGG dataset (See “Datasets” subsection). We construct a biological network to identify important pathways and related genes. Suppose 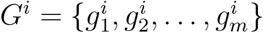 represents a set of genes in set *S* associated with *i*-th pathway *p_i_*. A biological network associated with pathways is considered as an undirectional weighted graph *G* =(*V, E, ω*). In graph *G*, each vertex represents a pathway. The two pathways *p_i_* and *p_j_* in graph *G* are connected by an edge *e_ij_* ∈ *E* whenever 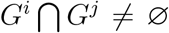. These two pathways are named neighbors, and the set of neighbors of each vertex is denoted by *N*(*p*). The *ω*(*e_ij_*), as the weight of the edge *e_ij_*, represents the number of genes in common between the two pathways *p_i_* and *p_j_*. For *H* ⊆ *V* we have 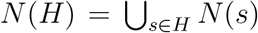. A path between two vertices *u* and *v* is a distinct sequence of graph vertices *u* = *v*_0_, *v*_1_,…, *v_k_* = *v* if each *v_i_v*_*i*+1_ (1 ≤ *i* ≤ *n* – 1) is an edge of graph *G*. The length of a path is equal to the number of edges in this path. The distance between two vertices *u* and *v* is equal to the length of the shortest path between these two vertices, which is denoted by *d*(*u,v*).

### C. The topological features of biological network

In this part, for each pathway, we define six informative features that correspond to the pathway’s position in the biological network:

1. The ratio of the number of genes in each pathway to the total number of COVID-19 related genes (*S*).

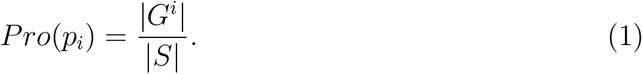 The larger value of this measure indicates that this pathway corresponds to the COVID-19 related genes.
2. The number of neighbors of each pathway in a biological network is called the degree of that pathway. The degree of each pathway is another topological feature that we have considered. A large value of this measure indicates the importance of the COVID-19 related genes in this pathway in terms of the number of participants in the other pathways.
3. Betweenness centrality value for each pathway, *p_i_*, in a weighted graph *G* is defined as follows:

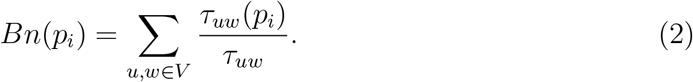 The value *τ_uw_* indicates the total number of shortest paths from *u* to *w* and *τ_uw_*(*p_i_*) shows the number of shortest paths that pass through *p_i_*. This measure shows the importance of the pathway *pi* in terms of centrality.
4. Let *L*(*G*) = [*l_ij_*] indicates the Laplacian matrix of weighted graph *G*, where:

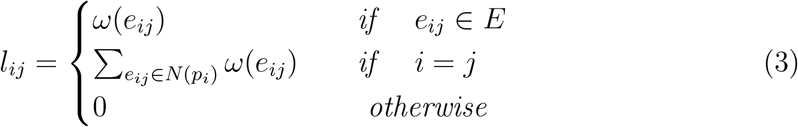 Assume that vector *u* = (*u*_1_, *u*_2_,…, *u_n_*) represents the normalized eigenvectors of matrix *L*(*G*) and vector (*λ*_1_, *λ*_2_,…, *λ_n_*) represents the associated eigenvalues [27]. Based on the second smallest eigenvalue, we compute the eigenvectors of the Laplacian matrix *L*(*G*) and store them in vector *X* = (*x*_1_,…, *x_n_*). Then, we sort the elements of vector *X* and place half of the nodes in partition *G*_1_ and the rest in partition *G*_2_. The subset *C* is selected from the vertices of two partitions *G*_1_ and *G*_2_ alternatively, such that removal of *C* separates the nods of graph *G* into two disjoint parts of almost equal size. Now the essentiality of each pathway, *p_i_*, is defined as follows:

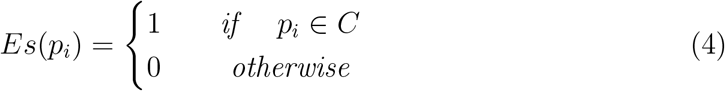
5. Let 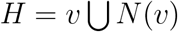, the Expansion of *H* is defined as follows:

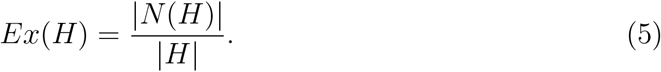

where |*N*(*H*)| and |*H*| demonstrate the size of *N*(*H*) and *H* respectively.
6. The Closeness centrality measure for each vertex *v* of graph *G* is defined as follows:

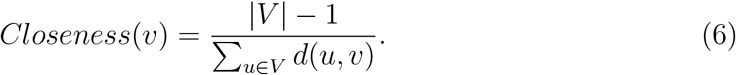 If the *Closeness*(*v*) value meets or exceeds a predefined threshold, we consider that the vertex *v* has the shortest distance with other vertices of the graph.

### D. Proposed model

Suppose *f_r_i__* represents the *r*-th feature of the *i*-th sample (1 ≤ *i* ≤ *n* and 1 ≤ *r* ≤ *m*). We assign a feature vector 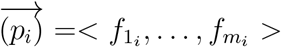 to each sample *p_i_* and define the column matrix *F_r_* = [*f*_*r*_1__,…, *f_r_n__*]^*T*^. To find the appropriate score for each feature we compute the following measures:

1. The weight matrix *S* = [*s_ij_*]_(*nn*)_ is defined as follows:

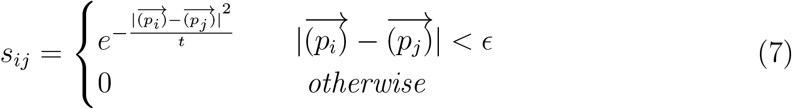
2. Suppose that *J* = [1, 1,…, 1]^*T*^ and let *L* = *D* – *S* indicates the Laplacian matrix then we define *D* = [*d_ij_*]_(*nn*)_ as follows:

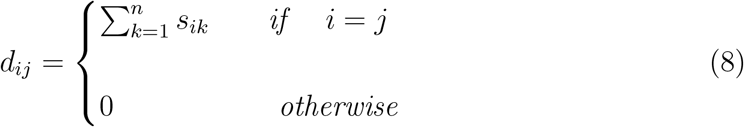
3. For the *r*-th feature, we define *F_r_* as follows:

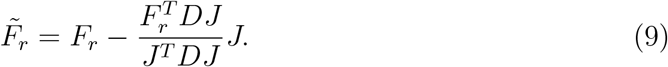
4. For the *r*-th feature, compute the Laplacian Score as follows:

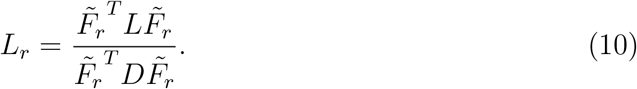

### E. Finding important COVID-19 related pathways

Since the problem of finding the appropriate set of COVID-19 related pathways is still an open question, it can be considered as a problem without a response variable or exact answer. Therefore, to find an efficient answer, we present a novel unsupervised learning method to find important COVID-19 related pathways. It is worth mentioning that, in supervised learning methods, feature selection has been studied extensively. Due to the lack of information about class labels to help the search for relevant knowledge in unsupervised learning methods, scoring features is a significantly more difficult challenge [28]. Our method comprises two main steps.

In the first step, we construct a biological network with respect to a set of COVID-19 related genes (S). Then we calculate six informative features for each pathway with respect to the topological features of the vertex in the biological network. It specifies a feature matrix *F* = [*f_ir_*] where *f_ir_* represents the *r*-th feature of the *i*-th pathway.

In the second step, we use an unsupervised feature selection method on a feature matrix to calculate the Laplacian Score for each feature (*L_r_*). Then, for the pathway *p_i_*, compute the Laplacian Score as follows:

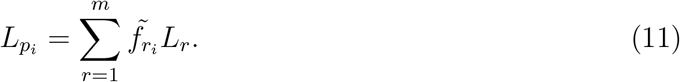

where 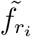 represents the normalized feature for the *i*-th pathway.

Finally, the high score pathways are selected as important COVID-19 related pathways.

## 5. Results

### A. Datasets

Identifying associated genes with disease pathology plays an important role in recognizing significantly related pathways and finding appropriate drugs. We collected four different sets of COVID-19 related genes. We utilized a set of experimental unapproved drugs used for COVID-19 treatment that are available on the Drug Bank [25]. This set is represented by Covid-Drug and includes 64 drugs. From these 64 drugs, 21 of them can target human proteins. These 21 drugs target 88 human proteins, which we defined as the first set of COVID-19 related genes and is denoted by the set *Co*. We also used a set of experimental unapproved drugs in clinical trials for COVID-19 treatment that are available on the Drug Bank [25]. This set includes 708 drugs, of which 347 drugs have been studied clinically in more than one clinic. From these 347 drugs, 213 drugs can target human proteins. This group of drugs is represented by the Clinical-Drug. These 213 drugs target 930 human proteins, which we defined as another set of COVID-19 related genes, denoted by the set *Cl*. The third set, which we considered as a set of COVID-19 related genes, contains 332 human proteins identified by Gordon et al. as targets of the virus in human proteins and is denoted by the set *T* [29]. The last set that we have considered as a set of COVID-19 related genes includes 93 human proteins presented by Habibi et al. is represented by set *E* [30]. Finally, we placed all of the above-mentioned four sets into set *S*. This set contains 1,300 genes, which we introduced as a final set of COVID-19 related genes. We also used the KEGG dataset, which contains complete information about pathways inside cells. From this dataset, we extracted information about signaling and disease pathways and related genes in each pathway. From the KEGG dataset, we selected two sets of signaling and disease pathways associated with COVID-19 that contain more than eight genes in the selected set *S*. In addition, to identify important genes in each pathway, we used the PPI network collected by Habibi et al [30]. This network contains 304,730 interactions between 20,040 human proteins. They collected information on the physical interactions between proteins in 5 datasets, including Biological General Repository for Interaction Datasets (BioGRID) [31], Agile Protein Interactomes Data analyzer (APID) [32], Homologous interactions (Hint) [33], Human Integrated Protein-Protein Interaction reference (HIPPIE) [34], and Huri [35]. They mapped all of the proteins from all of the above-mentioned five datasets to their corresponding Universal Protein Resource (UniProt) ID [36] and removed proteins if they could not be mapped to Uniprot IDs.

### B. Pre-processing

Since the spread of COVID-19, different techniques have been used to identify COVID-19 related genes. In this study, we used four different groups of COVID-19 related genes. Figure 3 shows the venn diagrams associated with these selected sets. Figure 3 demonstrates that more than 50% of the selected genes in each set, *Cl, E*, and *T*, are not in the other sets. In other words, these sets almost contain unique genes. In total, these sets include 1,300 genes associated with COVID-19, which we select as the COVID-19 related genes and denote by set *S*. Since many studies have been conducted to find out genes related to the pathology of COVID-19 and it is not possible to check all of them, we have reviewed a number of these studies. The results of our study show that the candidate genes in our selected set contain most of the important genes that are proposed through other studies. These studies used different methods and approaches for finding genes. In the following, we compare our candidate genes set with four different genes sets in other studies.

**Figure 3:**
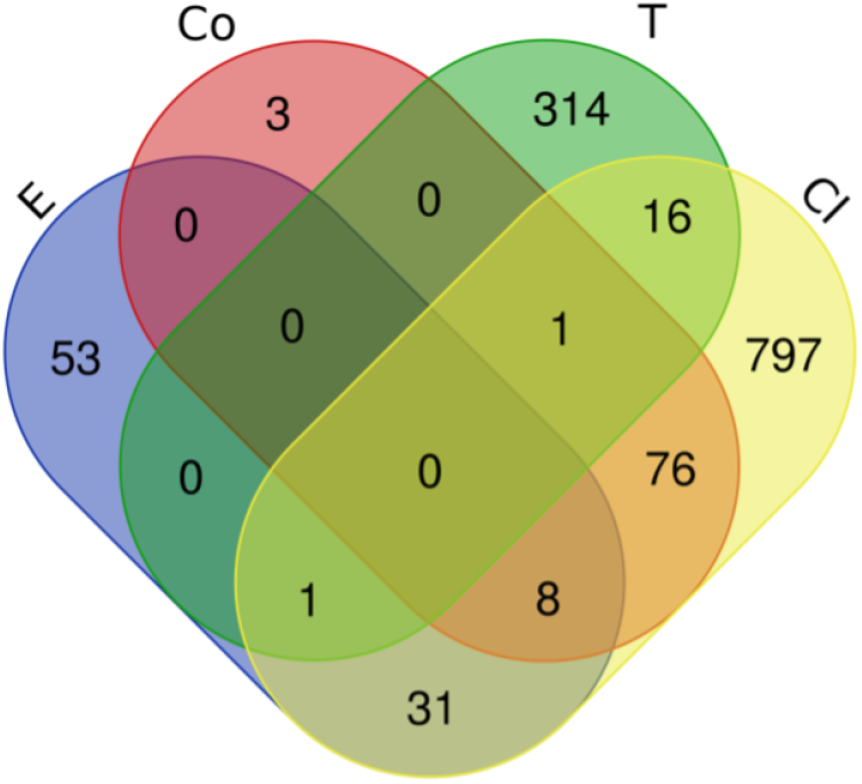
Venn diagram of COVID-19 related genes in sets E, Cl, Co, T respectively.

### C. Evaluation of COVID-19 selected sets of proteins

As mentioned earlier, we compared our selected gene sets with four gene sets that have been presented in other studies. The authors of the first study presented a method that is based on informative statistical and topological features in a human-virus protein interaction network [37]. More than 96% of the genes presented in [37] are represented by our candidate genes. The second study identified potential targets for repurposing based on Mendelian randomization [38]. Our candidate genes contain more than 95% of the genes presented in [38]. The authors of the third study investigated the COVID-19 comorbidity-associated gene sets [39]. Our candidate genes contain more than 84% of the genes presented in [39]. The last study identified the hub gene in the pathways related to COVID-19 [40]. Our candidate genes include more than 94% of the genes listed in [40]. Therefore, it can be concluded that our selected set *S* contains more than 92% of the genes reported in the above-mentioned studies. Thus, the selected set *S* could be a complete set of COVID-19 related genes.

We also studied six sets of disease-related genes for six underlying diseases: cardiovascular diseases, diabetes, hepatitis, lung diseases, kidney disease, and different types of cancers respectively. Then we compared the genes of these six sets with our selected set *S*. We found that from our 1,300 candidate genes, 461 genes are related to these six underlying diseases. From these 461 genes, 12 genes (APOE, CCR5, CD14, IL1A, IL1B, IL6, ITGB3, LDLR, MMP3, MTHFR, TNF) are commonly shared between all of the six disease-related genes. We also found 33 genes in more than 5 diseases out of 6 mentioned diseases. Figure 4 shows the percentage of these shared genes between disease.

**Figure 4:**
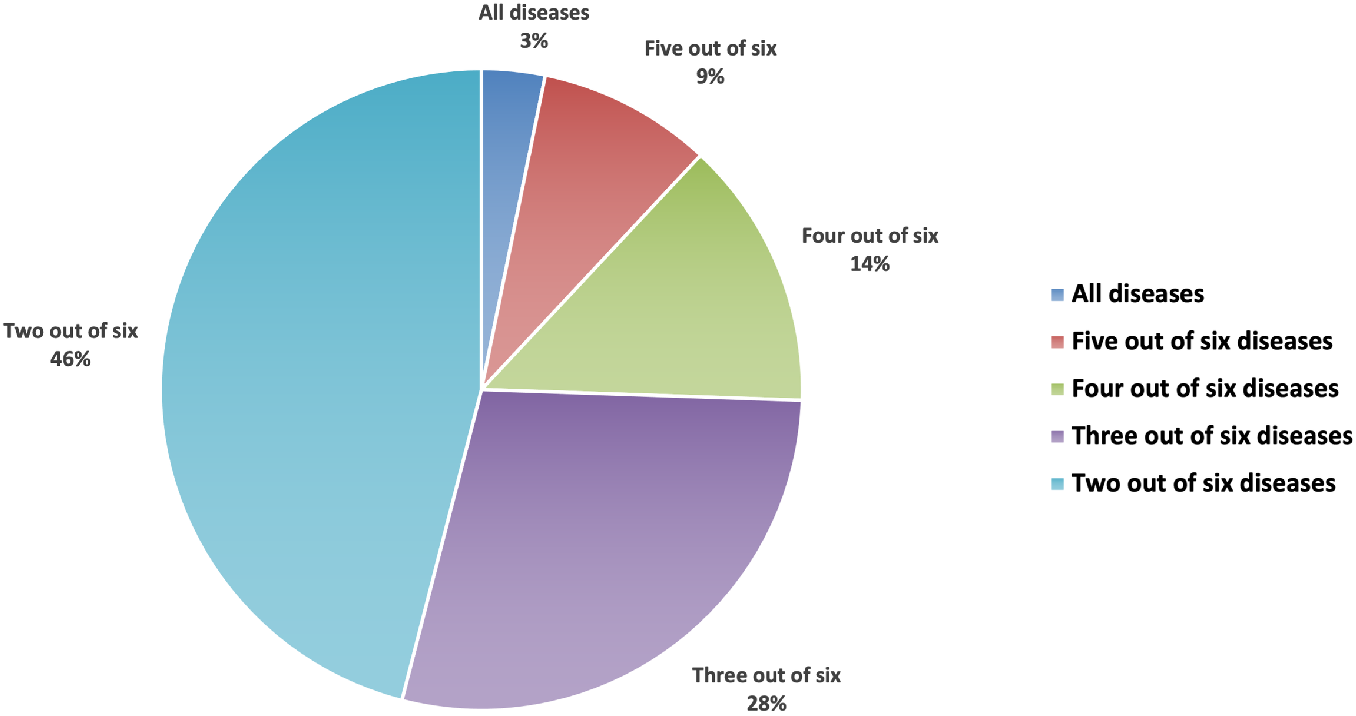
Percentage of shared genes in more than 2, 3, 4, 5 and 6 out of 6 underlying diseases.

### D. Evaluation of COVID-19 related pathways

One of the effective methods for finding therapeutic procedures in patients with severe COVID-19 is to understand the related molecular pathways. Identifying these pathways that are related to underlying COVID-19 could help to recognize potential drug targets.

Therefore, we studied the KEGG pathways associated with mentioned 1,300 COVID-19 related genes. For these 1,300 genes in *S*, there are 39 signaling pathways and 56 disease pathways. Table 1 presents the number of pathways related to our four candidate sets of COVID-19 related genes. Table 1 shows that the set of genes presented in set *S* covers the highest number of signaling pathways and the highest number of disease pathways as well. Most of these pathways have been studied as COVID-19 related pathways in other studies [41, 42, 43].

**Table 1:** The number of related pathways in four COVID-19 related selected gene sets.

|  | **Co** | **Cl** | **T** | **E** | **S** |
| --- | --- | --- | --- | --- | --- |
| No. Signaling pathway | 4 | 15 | 4 | 22 | 39 |
| No. Disease pathway | 12 | 23 | 2 | 31 | 56 |

Then, we build two biological networks associated with each set of signaling and disease pathways and calculate the informative features for each network. The informative features of these two biological networks are modeled by the Laplacian matrix. The output of the model is the Laplacian Score for each feature. Tables 2 shows the top ten signaling pathways with respect to the highest Laplacian Score and the number of genes in each pathway. Some other studies also introduced these pathways with other procedures and without measuring any priority as COVID-19 important related pathways. The last column of Table 2 shows the studies that presented these important pathways.

**Table 2:**
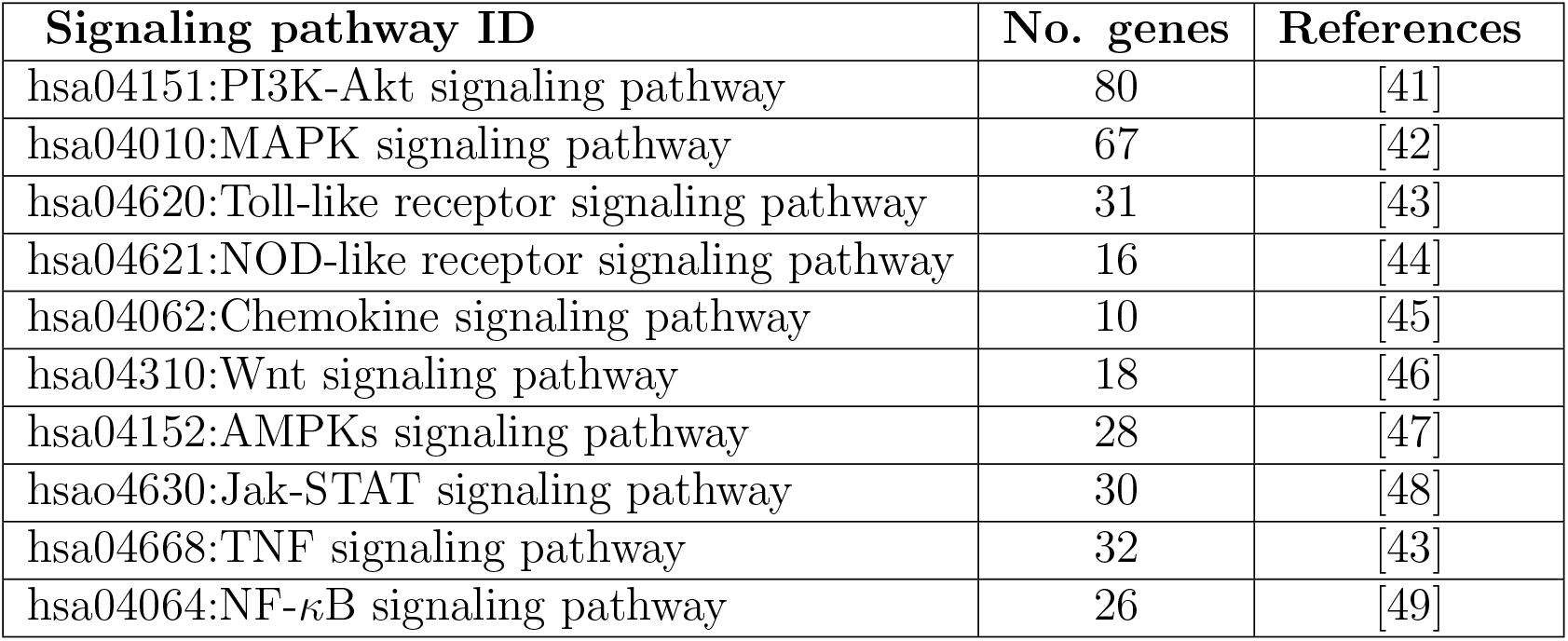
List of top 10 signaling pathways with respect to the highest Laplacian Score.

| **Signaling pathway ID** | **No. genes** | **References** |
| --- | --- | --- |
| hsa04151:PI3K-Akt signaling pathway | 80 | [41] |
| hsa04010:MAPK signaling pathway | 67 | [42] |
| hsa04620:Toll-like receptor signaling pathway | 31 | [43] |
| hsa04621:NOD-like receptor signaling pathway | 16 | [44] |
| hsa04062:Chemokine signaling pathway | 10 | [45] |
| hsa04310:Wnt signaling pathway | 18 | [46] |
| hsa04152:AMPKs signaling pathway | 28 | [47] |
| hsa04630:Jak-STAT signaling pathway | 30 | [48] |
| hsa04668:TNF signaling pathway | 32 | [43] |
| hsa04064:NF-κB signaling pathway | 26 | [49] |

Tables 3 also shows the top ten disease pathways with respect to the highest Laplacian Score and some of the important genes in each pathway.

**Table 3:** List of top 10 disease pathways with respect to the highest Laplacian Score.

| Disease pathway ID | No. genes | Important genes |
| --- | --- | --- |
| hsa05200:Pathways in cancer | 98 | P53, STAT3 |
| hsa05164:Influenza A | 56 | TNFRSF1A, IKBKB, JAK1 |
| hsa05166:HTLV-I infection | 49 | P53, MYC, JAK1 |
| hsa05152:Tuberculosis | 44 | TNFRSF1A, CASP3 |
| hsa04931:Insulin resistance | 34 | MTOR, IKBKB |
| hsa05169:Epstein-Barr virus infection | 30 | MYC, P53 |
| hsa05321:Inflammatory bowel disease (IBD) | 20 | IL2, STAT3 |
| hsa05323:Rheumatoid arthritis | 28 | CCL2, TNF, CCL5 |
| hsa05144:Malaria | 21 | TGFB1, CXCL8 |
| hsa05130:Pathogenic Escherichia coli infection | 10 | CD14, ACTB |

#### D.1. High score signaling pathways

In this subsection, we investigate three of the top high score signaling pathways that are selected through our method.

1. **PI3K/AKT signaling pathway** The first pathway is the Phosphatidylinositol 3-Kinase PI3K/AKT signaling pathway as one of the possible therapeutic targets for COVID-19 treatment. This pathway is associated with different characteristics of virus entry into the cell and the evolution of immune responses. PI3K/AKT signaling pathway performs a major role in cell invasion, growth, migration, and proliferation. It can inhibit apoptosis and increase angiogenesis. Irregular PI3K/AKT signaling pathway can produce diseases, such as cancer, and inhibitors of this pathway perform a crucial role in cancer treatment. On the other hand, it has been shown that SARS-CoV-2 endocytosis happens through a clathrin-mediated pathway which is regulated by the PI3K/ AKT signaling [41]. Therefore, suppression of this pathway can inhibit the entry of other viruses that use clathrin-mediated endocytosis. Other studies show that activation of the PI3K/AKT signaling pathway has been connected to the initiation of lung tissue fibrosis, which has been detected in COVID-19 patients. Based on this evidence, the use of anti-cancer and anti-viral drugs, to inhibit the PI3K/AKT signaling pathway and eventually suppress inflammation in COVID-19 can be an effective treatment for COVID-19 patients.
2. **MAPK signaling pathway** The second pathway is the p38 Mitogen-Activated Protein Kinase (MAPK). The p38 MAPK is the third major signaling cassette of the MAPK. This pathway performs a critical role in the release of pro-inflammatory cytokines like interleukin-6 (IL-6) and has been involved in myocardial dysfunction and acute lung injury. The strong inflammatory response in COVID-19 may be caused by abnormally upregulated p38 activity and can be explained by two mechanisms. First, during SARS-CoV-2 viral entry, angiotensin-converting enzyme 2 (ACE2) activity is lost. Second, as previously shown in SARS-CoV, p38 activity is upregulated via a viral protein, like other RNA respiratory viruses that may hijack p38 activity to promote replication [42]. Thus, SARS-CoV-2 may cause vast inflammation by directly activating p38 and down-regulating a key inhibitory pathway. The SARS-CoV-2 can simultaneously take advantage of p38 activity to replicate. Based on this evidence, p38 inhibitors should be considered for clinical trials in serious COVID-19 infections.
3. **TLRs signaling pathway** The next important pathway is the Toll-like Receptors (TLRs) as a family of ten members in humans. TLRs are expressed on different immune cells, like macrophages, dendritic cells, natural killer cells, B cells, and T cells. The innate immune system of this pathway is activated by many viruses, which is important for the removal of viruses. This procedure could be harmful to the host due to continuous inflammation and tissue damage. In patients with severe COVID-19, the inflammatory response begins with the release of cytokines, such as IL-6 and Tumoral Necrosis Factor-alpha (TNF-*α*). It is remarkable that these two main cytokines involved in severe COVID-19 (IL-6 and TNF-*α*) are downstream of TLR4. On the other hand, TRLs can activate the Janus kinase transducers (JAK/STAT), which could cause macrophage activation syndrome. Therefore, TLRs have a considerable dual role in viral infections [43]. Based on these evidences, TRLs properties make them possible candidates to treat inflammation in severe viral diseases like COVID-19.

Figure 5 shows that TLR signaling pathways have cross-talk with both PI3K/AKT and MAPK signaling pathways. TLRs are divided into two groups: a MyD88-dependent pathway and a MyD88-independent pathway. The first one drives the production of proinflammatory cytokines with the immediate activation of NF-*κ*B and MAPK. The second one associates with the induction of IFN-*β* and IFN-inducible and activation of NF-*κ*B and MAPK. In the MyD88-dependent pathway, the MyD88 protein recruits IRAK family proteins. The IRAK4 protein activates TRAF6 and this protein ultimately activates MAPKs and NF-*κ*B resulting in the production of proinflammatory cytokines. On the other hand, activated TRIF binds to TRAF3 and TBK1, ultimately activating IRF3 and IRF7 to initiate the production of interferons. TRIF also interacts with TRAF6 to stimulate MyD88-independent activation of MAPKs and NF-*κ*B. Our results in this subsection show the importance of TLRs in other mentioned signaling pathways. This relevance could confirm the potential link between the unknown COVID-19 pathways and the TLRs receptors.

**Figure 5:**
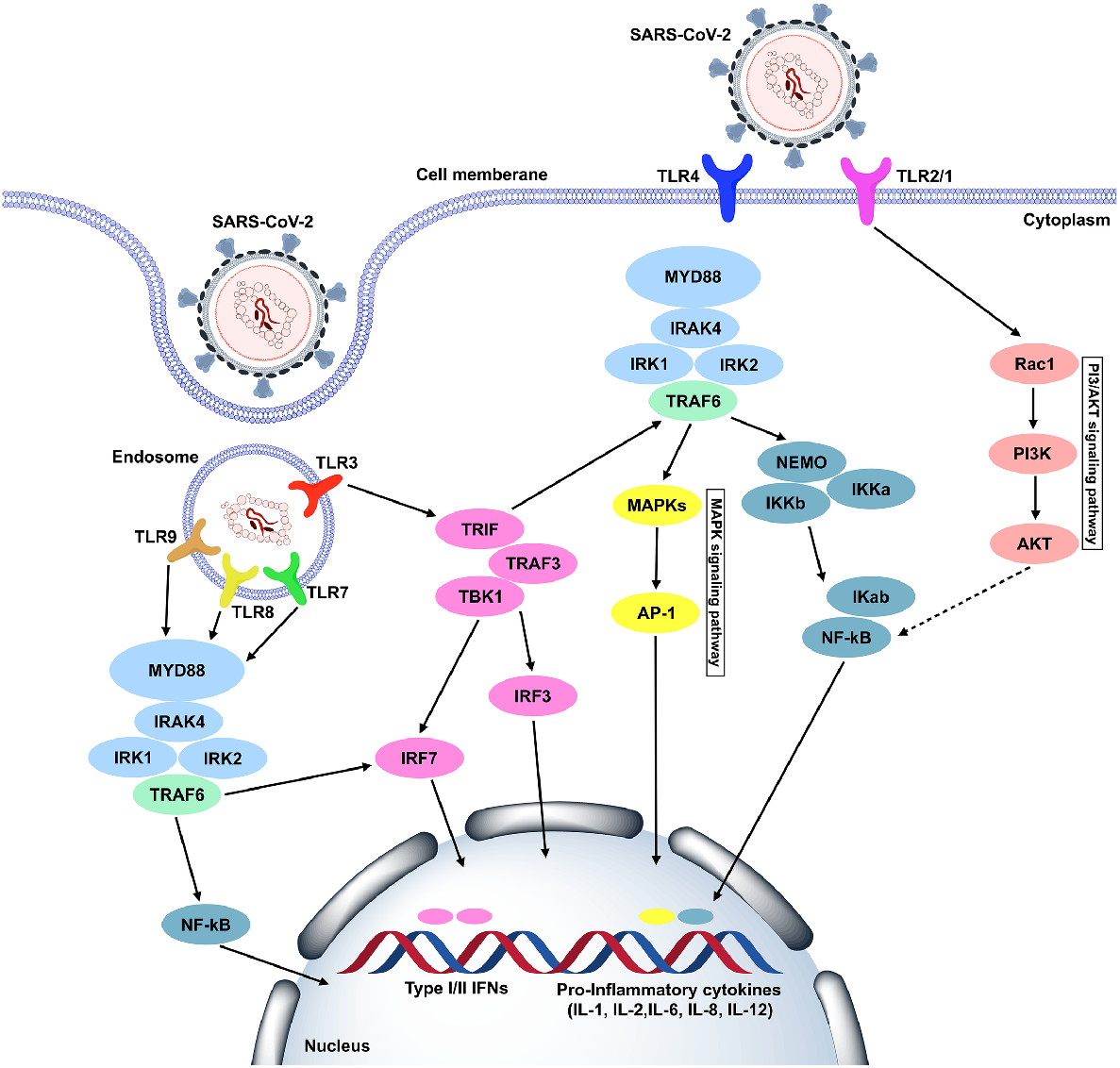
TLR signaling pathway cross-talks and immune deficiencies.

#### D.2. High score disease pathways

In this subsection, we discuss three of the most important disease pathways and related important genes that have the highest score through our method. Identifying the shared pathways between underlying diseases and COVID-19 and also identifying the important genes related to each pathway gives us valuable information about possible drug targets.

1. **Pathways in cancer** The first COVID-19 related disease pathway is the pathways in cancer (*hsa*05200). We find that 98 genes are shared between pathways in cancer and COVID-19. Figure 6 shows the PPI network between these 98 genes. We evaluate two important genes TP53 and STAT3 with the valuable topological properties in this network. In Figure 6 the topologically important genes are shown in red.
  a. **TP53**: The first important gene that is shared between pathways in cancer and COVID-19 is the Tumor Protein P53 (TP53 or P53). The cellular tumor antigen P53 is a key member in INF type 1 [50]. Researchers believe that the key tumorsuppressor P53 protein will undergo degradation by SARS-CoV-2, leading to increased viral survival in host cells. Thus, P53 activators might effectively inhibit viral replication in the human respiratory tract and lung cells [51]. Therefore, TP53 could be a potential therapeutic target for COVID-19 treatment.
  b. **STAT3**: The second important gene that is shared between pathways cancer and COVID-19 is the Signal Transducer and Activator of Transcription 3 (STAT3). Recent studies showed that the viral components of SARS-CoV-2 cause the abnormal hyperactivation of STAT3 through COVID-19 cytokines [52]. Abnormal activity of STAT3 elevates lymphopenia and reduces anti-virus immune responses. Therefore, STAT3 could be a potential therapeutic target for COVID-19 treatment.
2. **Influenza A pathway** The second one is the Influenza A pathway (*hsa*05164), which is identified as an essential disease pathway related to COVID-19. We find that 57 genes are shared between Influenza A and COVID-19. We analyze the topologically important genes among these 57 reported genes. This analysis is accomplished with respect to the proteinprotein interaction (PPI) network. As shown in Figure 7, three genes, TNFRSF1A, JAK1, and IKBKB, are known as important genes. In Figure 7 the topologically important genes are shown in red.
  a. **TNFRSF1A**: The first important gene that is shared between Influenza A and COVID-19 is TNFRSF1A/TNFR1. The Tumor Necrosis Factor Receptor (TNFRs) is recognized as the receptor for TNFSF2/TNF-*α* [36]. TNF and its receptors (TNFR1 and TNFR2) are related to proinflammatory responses and play a major role in the regulation of inflammation. Previous studies have shown the relation between the severity of COVID-19 and increasing TNFR1 [53]. Therefore, TNFRSF1A could be a potential therapeutic target for COVID-19 treatment.
  b. **JAK1**: The second important gene that is shared between COVID-19 and Influenza A is the Tyrosine-protein kinase JAK1 (JAK1). JAKs (JAK1 and JAK2) are transmembrane proteins that increase extracellular signals. Many of the cytokines, including interferons (IFNs), involved in the cytokine storm use protein kinases JAKs for signal transduction [48]. Therefore, blocking IFNs which are mediated by JAK could be an effective COVID-19 treatment.
  c. **IKBKB**: The third important gene that is shared between between Influenza A and COVID-19 is the Inhibitor of nuclear factor *κ*B kinase subunit beta (IKBKB or IKK2). IKBKB plays an essential role in the transcription factor nuclear factor-*κ*B (NF-*κ*B) pathway. This pathway changes multiple pathogenic activities such as inflammatory cytokines [49]. Several studies confirmed the benefit of IKKs as a downstream effector of this pathway to weaken COVID-19. Therefore, IKBKB could be a potential therapeutic target for COVID-19 treatment.
3. **HTLV-I infection pathway** The third COVID-19 related disease pathway is the HTLV-I infection pathway (*hsa*05166). We find that 49 genes are shared between the HTLV-I infection pathway and COVID-19. Figure 8 shows the protein interaction network between these genes. We evaluate three important genes, MYC, P53, and JAK1 with the valuable topological properties in this network. In Figure 8 the topologically important genes are shown in red.
  a. **MYC**: In the previous sections, we discussed P53 and JAK1. The next important gene that is shared between the HTLV-I infection pathway and COVID-19 is the Myc proto-oncogene protein (MYC). Recent studies have shown an increase level of MYC in COVID-19 patients. The regulation of MYC may contribute to cell proliferation, especially macrophages and neutrophils [54]. Therefore, MYC could be a potential therapeutic target for COVID-19 treatment.

**Figure 6:**
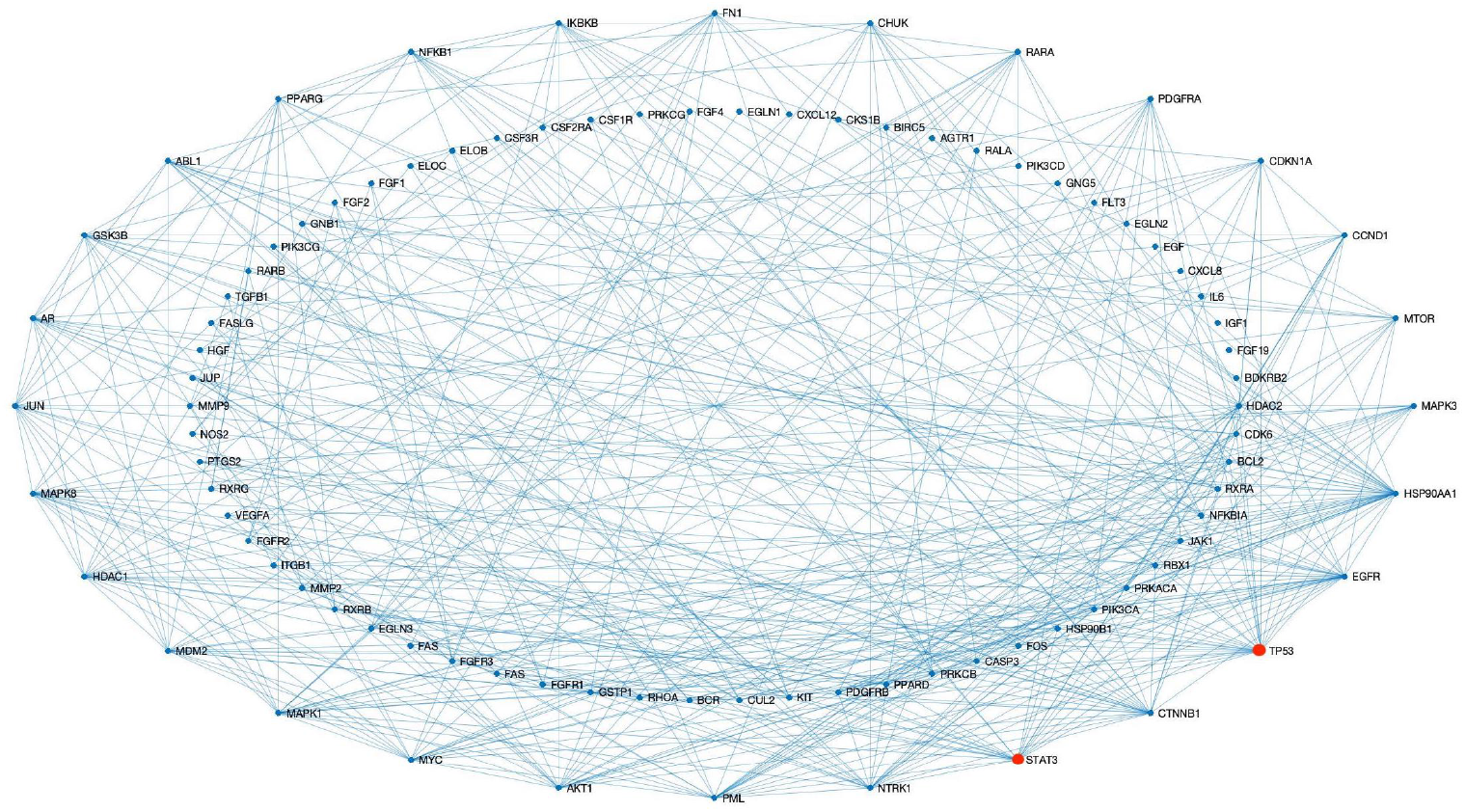
The PPI network associated with 98 genes are shared between pathways in cancer and COVID-19.

**Figure 7:**
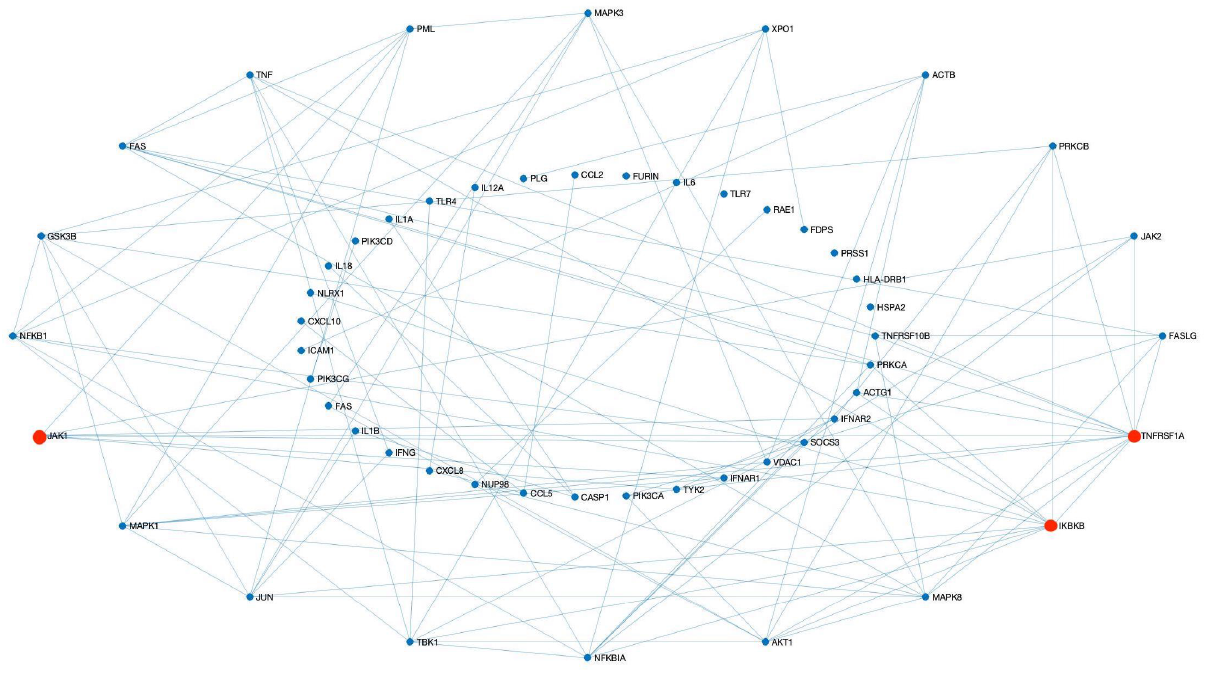
The PPI network associated with 56 genes are shared between Influenza A and COVID-19.

**Figure 8:**
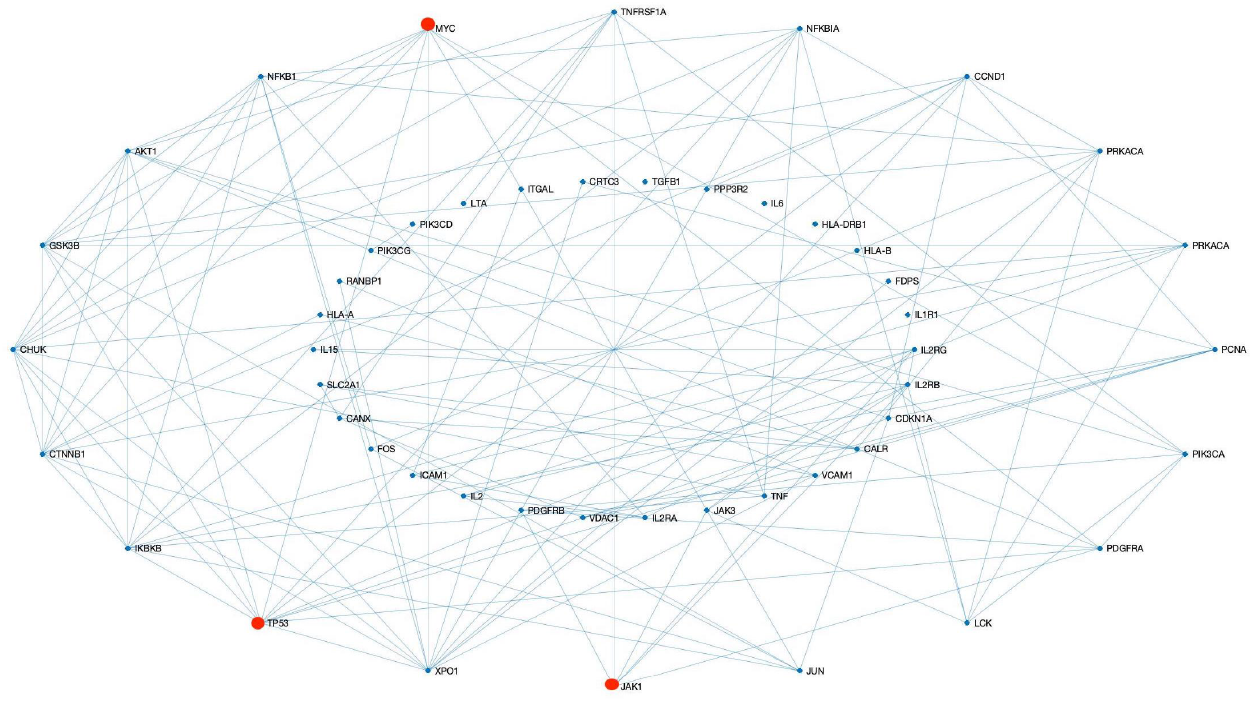
The PPI network associated with 49 genes are shared between the HTLV-I infection pathway and COVID-19.

We also compare and evaluate the COVID-19 related genes for three highly scored disease pathways. Figure 9 shows the venn diagrams associated with these disease pathways. Figure 9 demonstrates that 11 genes were identified through all three disease pathways. Table 4 shows the complete list of these 11 genes and detailed information about their corresponding functions.

**Figure 9:**
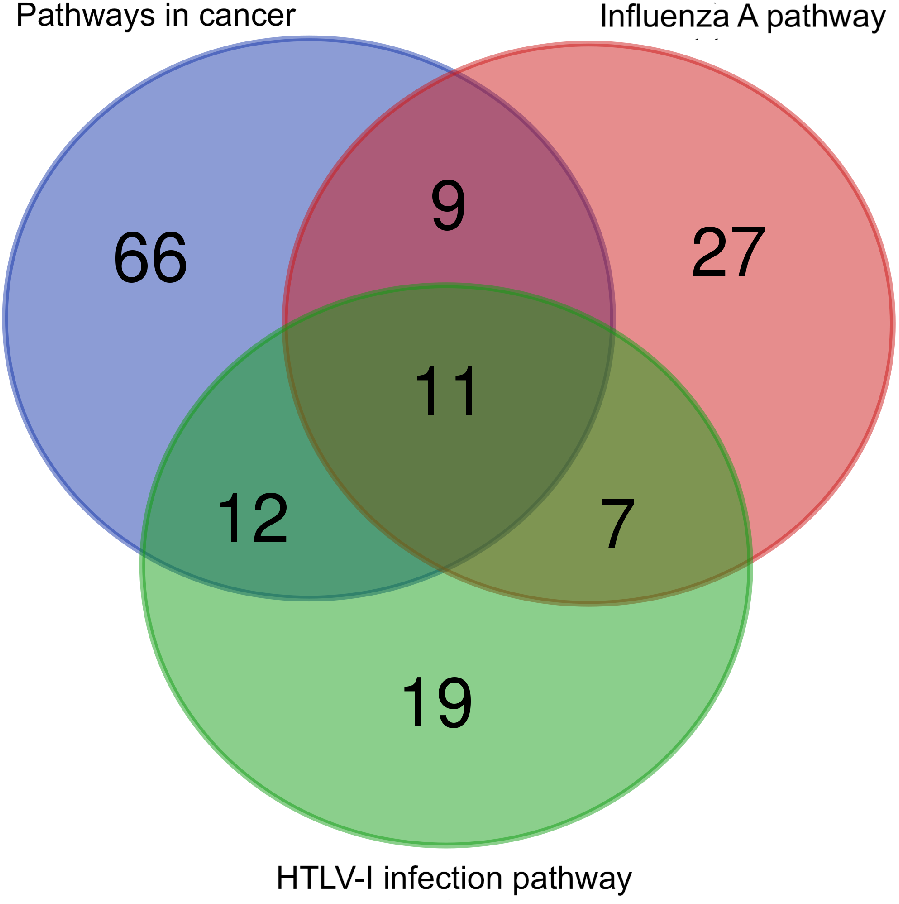
Venn diagram of COVID-19 related genes in three highly scored disease pathways.

**Table 4:** The list of shared genes between three disease pathways and their functions.

| Gene name | Function |
| --- | --- |
| NFKBIA | The NFKBIA helps keep NF-κB bound in the IKK complex. NF-κB regulates the activity of multiple genes, including genes that control the body's immune responses and inflammatory responses [55]. |
| PIK3CD | The PIK3CD provides instructions for an enzyme called phosphatidylinositol 3-kinase (PI3K). PI3K is found in white blood cells, including immune system B cells and T cells. These cells identify and attack foreign invaders like viruses and bacteria and prevent infection [55]. |
| JUN | The JUN encodes a protein that is highly similar to the viral protein. This gene interacts directly with precise target DNA sequences to control gene expression. JUN is mapped to a chromosomal region involved in both translocations and deletions in human malignancies [55]. |
| JAK1 | The JAK1 has a critical role in influencing the expression of genes that mediate inflammation. This gene is an important part of the (IL-6)/JAK1/STAT3 inflammation and immune response, and it could be an effective drug target for reducing cytokine storms [55]. |
| AKT1 | The AKT1 encodes one of the three members of the human AKT serine-threonine protein kinase family. AKT proteins control a large number of cellular functions like cell proliferation, metabolism, survival, and angiogenesis in both malignant and normal cells [55]. |
| GSK3B | The GSK3B encodes a protein that belongs to the glycogen synthase kinase subfamily. It is involved in energy metabolism, inflammation, ER-stress, mitochondrial dysfunction, and apoptotic pathways [55]. |
| IL6 | The IL6 encodes a cytokine that operates in inflammation and in the maturation of B cells. The function of IL6 is implicated in a wide range of inflammation-associated diseases. High levels of the encoded protein have been found in virus infections like COVID-19 [55]. |
| PIK3CG | The PIK3CG encodes a protein that is a class I catalytic subunit of PI3K. Since PI3K is found in the immune system, which identifies and attacks viruses and bacteria, it prevents infection. |
| NFKB1 | The NFKB is a transcription regulator that is activated by various cellular stimuli like cytokines, bacterial and viral products. It has a major role in the regulation of the early response to viral infection. Inappropriate activation of NFKB has been associated with several inflammatory diseases [55]. |
| IKBKB | The IKBKB encodes a protein that causes dissociation of the inhibitor and activation of NF-κB which is activated by multiple stimuli such as inflammatory cytokines, bacterial or viral products [55]. |
| PIK3CA | The PIK3CA provides instructions for PI3K. Since PI3K is found in the immune system, which identifies and attacks viruses and bacteria, it prevents infection [55]. |

## 6. Conclusion and Discussion

There are knowledge gaps in the systems biology of COVID-19, and this makes the treatment of this disease more difficult. Continuous and comprehensive research with different points of view tries to identify these gaps and eliminate these shortcomings. Recently, many studies have been presented to identify common pathways between COVID-19 and other known signaling and disease pathways. In these studies, the potential relationship between COVID-19 and other diseases has been discussed. Therefore, a large number of signaling and disease pathways related to COVID-19 have been explored. Recognizing a smaller number of more related and important pathways can help researchers conduct more investigations. For this purpose, we constructed two biological networks corresponding to signaling and disease pathways and defined some informative features. Then, we proposed an unsupervised method for ranking these important COVID-19 related signaling and disease pathways. In the first step of this work, we selected a set of genes associated with COVID-19. This selected set contains most of the genes that have been presented as drug targets for COVID-19 in the Drug Bank or have been reported in multiple studies as COVID-19 related genes. Then, we extracted the corresponding pathways from KEGG for these selected genes. Each of the extracted pathways contains at least eight genes from the candidate gene set. Afterward, we constructed the two biological networks corresponding to these COVID-19 related signaling and disease pathways. We also defined six informative features for these networks. Then, we used an unsupervised learning method to score each feature and sort pathways for each network with respect to the corresponding score.

The results of our proposed model on a biological network corresponding to signaling pathways showed that three signaling pathways, PI3K/AKT, MAPK, and TLRs, were selected through our method as the top three signaling pathways. These pathways have the most cross-talk with COVID-19. Some researches on TLRs signaling pathways showed that MyD88 related pathways activate NF-*κ*B and MAPK signaling pathways rapidly, and these pathways produce proinflammatory cytokines. Other studies also showed the cross-talk between TLRs and PI3K/AKT signaling pathways. Figure 5 shows the cross-talk between TLR signaling pathways and both PI3K/AKT and MAPK signaling pathways as well. Tolllike receptors (TLRs), as a family of ten members in humans, play an important role in identifying invasive pathogens and establishing an innate and adaptive connection. These receptors are expressed in immune cells and some other cells, but their regulation and expression in cells are associated with specific molecules derived from pathogens or damaged cells. The binding of the ligands to the TLRs activates the signaling cascade and initiates the body’s innate immune response. This immune response of cells leads to the production of cytokine proteins that can limit the spread of the virus in the body. But some inflammatory cytokines such as TNF-*α*, IFNs, and IL6 are also stimulated and produced. Failure to regulate these inflammatory responses in the immune system and a sudden increase in cytokine production can lead to loss of lung function, shock, and even death in patients with COVID-19. Therefore, our results indicate the importance of TLRs in all other mentioned signaling pathways, which could demonstrate the potential link between the unknown COVID-19 pathways and the TLRs receptors. Several trials are being conducted regarding TLRs’ pathways in COVID-19 that could lead to the finding of new drugs for COVID-19 treatment.

The results of our proposed model on a biological network corresponding to disease pathways showed that three disease pathways, pathways in cancer, Influenza A, and HTLV-I infection, were selected through our method as the top three disease pathways. Some of the important genes from these pathways are TNFR1, JAK1, IKK2, TP53, STAT3, and MYC, respectively. Recent studies have shown that these topologically important genes from our top disease pathways can be targeted by COVID-19 candidate drugs such as Baricitinib, Ruxolitinib, Tofacitinib, and Acetylcysteine, which are already approved or are undergoing clinical trials [25]. This could confirm that our method selected some important disease pathways and related genes. It can be concluded that 11 common and important genes that are selected through our method in all three disease pathways (Table 4) could be good candidates as drug targets for further research in clinical trials for possible COVID-19 treatment.

